# Large-scale neuron cell classification of single-channel and multi-channel extracellular recordings in the anterior lateral motor cortex

**DOI:** 10.1101/445700

**Authors:** Rohan Parikh

**Author notes:** The author has declared that no competing interests exist. Present address: 19700 Helix Drive, Ashburn VA, 20147.

## Abstract

Identification of neuron cell type helps us connect neural circuitry and behavior; greater specificity in cell type and subtype classification provides a clearer picture of specific relationships between the brain and behavior. With the advent of high-density probes, large-scale neuron classification is needed, as typical extracellular recordings are identity-blind to the neurons they record. Current methods for identification of neurons include optogenetic tagging and intracellular recordings, but are limited in that they are expensive, time-consuming, and have a limited scope. Therefore, a more automated, real-time method is needed for large-scale neuron identification. Data from two recordings was incorporated into this research; the single-channel recording included data from three neuron types in the motor cortex: FS, IT, and PT neurons. The multi-channel recording contained data from two neuron subtypes also in the motor cortex: PT_L and PT_U neurons. This allowed for an examination of both general neuron classification and more specific subtype classification, which was done via artificial neural networks (ANNs) and machine learning (ML) algorithms. For the single-channel neuron classification, the ANNs achieved 91% accuracy, while the ML algorithms achieved 98% accuracy, using the raw electrical waveform. The multi-channel classification, which was significantly more difficult due to the similarity between the neuron types, yielded an ineffective ANN, reaching 68% accuracy, while the ML algorithms reached 81% using 8 calculated features from the waveform. Thus, to distinguish between different neuron cell types and subtypes in the motor cortex, both ANNs and specific ML algorithms can facilitate rapid and accurate near real-time large-scale classification.

---

Brain and neuron activity may be visualized in two ways; observing neuron activity optically is useful, but its scope is limited solely to the surface of the brain. However, silicon probes have emerged as a more effective way of observing brain activity, especially the recently pioneered Neuropixel probe (1).

This activity correlates to particular neurons, specifically in the anterior lateral motor cortex for this research, which are classified as regular spiking (RS) and fast spiking (FS). RS neurons can be further broken down into excitatory putative pyramidal tract (PT) neurons and inhibitory layer 5 intratelencephalic (IT) neurons; meanwhile, FS GABAergic neurons are inhibitory (2). These L5 PT neurons in the motor cortex may be subdivided into PT_L & PT_U neurons, further subtypes in which preparatory activity to motor commands has been observed to indicate these neurons as having specialized distinct roles in motor control.

Currently, the electrophysiological data generated using Neuropixel probes from neuron cells is simply the general background electrical activity of the brain with occasional spikes from neurons firing. These spike waveforms have unique characteristics derived from the specific neurons they originate from, which are used to sort these spikes into various clusters. This clustering has been attempted using a swath of different techniques, including thresholding, feature extraction, and template-matching, and frequently require human correction due a semi-automated methodology. These issues are prevalent in nearly all spike sorting techniques that have been used on tetrodes and smaller-scale electrode arrays, along with those that have been built specifically for large-scale dense electrode arrays. These include Kilosort (3), Klusta (4), JRClust (5), M-Sorter (6), YASS (7), MountainSort (8), SpyKING CIRCUS (9), FAST (10), and various other unnamed algorithmic methods (11). Many of these algorithms claim full automation, but in reality require human correction to some degree (12). Thus, in order to accurately classify neurons, biological imaging, sequencing techniques, or intracellular recording is generally required. Optogenetic tagging is a bio-imaging technique in which a firing neuron is tagged in order to identify it. While this process is useful in identifying the specific neuron type responsible for firing to perform a behavior, it is time-consuming and is limited to single neurons, making it infeasible for large-scale classification (13). Single-cell RNA-sequencing using comprehensive transcriptome analysis is another biological technique to classify neuron types, but again, it is time-consuming and requires individualized analysis of single neurons (14). Neuron classification of hypothalamic supraoptic neurons in rats has used an electrophysiological approach through firing patterns, but also generally requires immuno-chemical labeling unless the patterns are in phasic bursting (15).

### Significance Statement

Identification of neuron type helps understand the connection between neural circuitry and behavior. With the advent of high-density probes, large-scale neuron classification is needed, as typical extracellular recordings are identity-blind to the neurons they record. The purpose of this research was to determine the viability of neuron classification via artificial neural networks and machine learning, and evaluate the hyperparameter and feature selection for such classification. The study yielded specific hyperparameters and features for certain algorithms and networks that consistently provide extremely high accuracies to serve as a basis for further classification without excessive optotagging and intracellular recordings.

Intracellular recording utilizes a patch clamp to measure the electrical activity within one neuron, providing the ground-truth data between spike waveform and cell type; however, this is limited to singular neurons and is not representative of the surrounding electrical activity. In comparison, extracellular recordings allow for the recording of many neuron cells firing; however, this introduces the trade-off in which spikes can be clustered but not identified (16). Accurate neuron classification into three classes of mouse cortical neurons and rat dorsal root ganglia has been achieved using intracellular recordings, and classification into four classes of cat primary visual cortical neurons has also been achieved with intracellular recordings, but these methods do not account for interference of electrical activity in the brain, seen in extracellular recordings, and is not automated due to non-software-based clustering such as parameter extraction (17) (18) (19). Current methods have also utilized both RNA-seq and single-cell patch-clamp (intracellular) protocols to identify neuronal subtypes, but also have a limited scope (20).

Other waveform classification algorithms may also be algorithmic and software-based, but have their own limitations. One method utilized manual K-means clustering to perform real-time classification, but was limited in that it solely used 8 time points in the waveform and was limited to 30 electrodes. Manual algorithms that are now automated with machine learning are not as effective, especially considering the large scale of current probes, with waveforms containing over 30 time points and 384 electrodes simultaneously recording (21) (22). Another method utilized a probabilistic approach through a Gaussian Process Classifier with a variational Bayesian approach and radial basis function, and achieved 72.5% to 92.7% accuracy in the univariate classification and up to 99.2% accuracy in twin-variate classification for several rat and cat cells. However, the accuracy ranged widely in different methods and was only performed for 40-120 neurons, which is not necessarily sufficient as justification for large-scale classification (23).

Research using neural networks in the classification of four types of adult human dentate nucleus neurons saw a mis-classification rate of 32.8% to 37.2% using topological data, and a misclassification rate of just 3.3% using morphological data; while useful, this data is significantly harder to obtain than electrophysiological data, which merely requires a probe with electrodes inserted into the brain (24). In addition, general classification of several myenteric neuron types has been shown to require morphological supplementary data to assist electrophysiological data in classification (25).

Thus, it seen that there are several pressing issues with regard to neuron cell classification. Spike sorting is plausible in conjunction with other identification methods such as optotagging or RNA-seq, but this requires repeated iterations of these techniques to confirm cluster identification. In addition, clustering methods that use spike sorting algorithms are time-consuming, which hinders real-time classification. The accuracy of these spike sorting algorithms is often variable and resulting clusters can be difficult to distinguish, because the algorithm will not definitively assign a cluster or classification to each spike waveform. Finally, artificial neural networks as a classification tool are promising, but require or recommend morphological data in conjunction to electrophysiological data.

Currently, neuron classification has been attempted and has seen success with extracellular recordings, both single-channel and multi-channel, in various brain regions including the primary visual cortex, cortical visual area AM, cortical visual area RL, hippocampus, lateral geniculate nucleus, lateral posterior nucleus, superior colliculus, and cerebellum. The classification techniques used in classification of neurons from these brain regions includes random forests, K-means clustering, and t-distributed stochastic neighbor embedding (t-SNE) (13). However, neuron cell classification has not yet been attempted in the anterior lateral motor cortex; in addition, artificial neural networks and various other promising machine learning algorithms have not been examined.

The purpose of this research is to develop an accurate neuron classification method for the anterior lateral motor cortex with single-channel and multi-channel electrophysiological extracellular recordings via multilayer perceptron neural networks (MPNs), convolutional neural networks (CNN), random forests (RF), K-means clustering, t-SNE, k-nearest neighbors (KNN), gradient tree boosting (GTB), extra trees (ET), and logistic regression (LR) classification. In the single-channel recordings, the purpose is to distinguish between distinct cell types, while in the multi-channel recording, the purpose is to distinguish subtypes of a specific cell. In doing this, the effects of classification metrics and hyperparameter tuning on accuracy is investigated as well.

## Materials and Methods

### Single-channel recording

Single-channel electrophysiological data was the alm-1 dataset obtained from CRCNS (26). Data preprocessing was performed in MatLab R2018 on the spike waveforms, which were a set of 29 single points that made a waveform when plotted. The L5 IT and PT cells in this dataset were optogenetically tagged with CRE-dependent AAV virus expressing ChR2, ensuring its ground-truth validity and verifying the set of mathematical analyses. The FS neurons were determined by a spike-sorting methodology due to the distinctly small time interval between spikes. Despite the lack of optogenetic tagging for these neurons, their identity is still known due to the unambiguous nature - in terms of fast-spiking compared to regular-firing - of GABAergic neuron spiking. Feature extraction was done as the waveforms were separated by cell type (cell types were FS, PT, & IT); eight features were calculated (See Appendix A.1 for full code).

#### Feature extraction & pre-processing

Figure 1 shows the distinct differences between three cell types found in the motor cortex; however, while they are visually very different, a feature extraction methodology is needed to quantify these waveforms to find mathematical patterns.

**Fig. 1.**
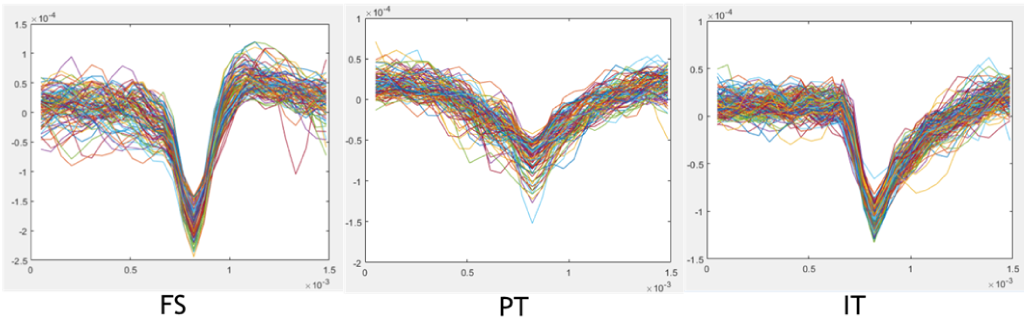
150 waveforms of 29 points superimposed to distinguish the waveform shape between the FS, PT, and IT neuron cell type.

The first three features were related to the waveform’ s amplitude. The first was the full amplitude (fA); it was calculated by the absolute difference between the peak and trough of the waveform (Appendix A.1 Line # 183-187). The second feature was the negative amplitude (nA), calculated as the difference between the trough and 0 (Appendix A.1 Line # 190-193). The third feature was, on the other hand, the positive amplitude (pA), calculated as the difference between the peak and 0 (Appendix A.1 Line # 195-199).

The fourth and fifth features were related to the width of the waveform. The fourth was the distance from the trough to the first peak following the trough, and was labeled the “recovery time” (rT; Appendix A.1 Line # 202-209). The fifth, alternatively, was the distance from the first peak in the waveform to the trough, and was labeled the “spike time” (sT; Appendix A.1 Line # 212-219).

The final three features were related to the group of waveforms within one trial. The sixth feature was the interspike interval (isi), calculated as the time difference between consecutive spikes (Appendix A.1 Line # 222-234). The seventh feature was the regularity of the spikes (reg), calculated as the variance of the ratio between consecutive interspike intervals (Appendix A.1 Line # 237-244). The eighth feature was burstiness (b), which was calculated as the number of interspike intervals that were less than a tenth of the mean interspike interval for a cell type, meaning the cell fired as a burst (Appendix A.1 Line # 248-257).

In addition to these eight calculated features, the entire waveform of 29 units was appended to form the final 29 features. Before these 29 units were added, the electrical background activity of the brain, or the noise, was base-lined to prevent it interfering with the classifiers (Appendix A.1 Line # 123-126).

After this preprocessing, the result was three separate matrices (FS, PT, IT) with n rows, with n equal to the number of waveforms for a given cell type, and 37 columns, one for each feature. These matrices were transferred to a Python IDE (Jupyter Notebook) for further processing and classification via a.csv file intermediary.

#### Training - testing set creation

Further processing was performed in Python 3 to create training and testing arrays for the future ANNs and ML algorithms (See Appendix B for full Python code of data processing and train/test set creation). The training data matrix was first filled with the existing data available for each neuron cell type (67% of the available data for each cell type was used in accordance with a 67:33 train-to-test split ratio; Appendix B Line # 98-105). Since there was a large discrepancy in the available data for the individual cell types (1,438,775 FS waveforms compared to 319,484 and 126,460 PT & IT waveforms), the ANNs and other ML algorithms may have only predicted the FS class for all waveforms. Thus, oversampling was performed to synthesize new data and equalize the training data for each neuron cell type. (Appendix B Line # 129-141).

After this, all neuron cell types were randomly sorted into master training and testing sets after NaNs were removed(Appendix B Line # 144-186). The 29 individual points of the waveform were extremely small values, on the order of 1e^−4^ and 1e^−5^; the discrepancy between these values and the calculated features would interfere with the classifier, so they were normalized to values between zero and one (this normalization was performed on the three amplitude calculations as well; Appendix B Line # 196-197).

### Multilayer perceptron neural network

The MPN was then trained and tested using PyTorch (See Appendix C for full Python code of MPN training and testing). Its architecture consisted of 3 hidden layers, with between 3-37 input nodes, depending on the feature selection of the trial, and 3 output nodes for the three classes. Each fully connected layer but the final one was followed by a rectified linear unit activation function (ReLU) as well; a log softmax activation function was performed on the final layer (Appendix C Line # 50-70). The feature selection variable was informed by the recursive feature elimination (RFE) algorithm. The set of the best three features (fA, nA, pA), best four features (fA, nA, pA, reg), and best five features (fA, nA, pA, reg, isi), along with all eight features, the 29 points of the waveform, and all 37 features together (Appendix C Line # 29-41) were selected. The learning rate was set at 0.001, the optimizer function was stochastic gradient descent with momentum (p = 0.9), and the loss function was CrossEntropyLoss (Appendix C Line # 76-79). Batch size was set at 100 (Appendix C Line # 101). Epoch number was variable to determine the minimum training needed to plateau accuracy and evaluate the speed at which the network learned; it was tested at 1, 2, 3, 5, 10, 25, 50, and 100 epochs.

### Convolutional neural network

The CNN was created and tested in PyTorch as well (See Appendix D for full Python code of CNN training and testing). Its network architecture consisted of 2 1-D convolutions, each of which was followed by a ReLU activation function and a 1-D pooling layer, and 2 fully-connected hidden layers. Each fully-connected hidden layer was followed by a ReLU activation function, and the final layer was followed by a log softmax activation function (Appendix D Line # 50-81). The feature selection, learning rate, optimizer function, loss function, and batch size were identical to the MPN. In addition, epoch number was varied identically.

### Machine learning algorithms

Seven additional ML algorithms were tested; three that were performed in existing literature in eight other regions of the brain (RF, k-means, and t-SNE). Four additional ones (KNN, GTB, ET, & LR) were promising and were tested as well (See Appendix E for full Python code of additional ML algorithms; all additional ML algorithms used Scikit-learn in Python).

The random forests algorithm, essentially swarm intelligence with decision trees, was performed. The number of decision trees was varied; 100, 500, and 1000 were tested. In addition, the number of maximum features were varied as well; 3, 4, 5, and 8 randomized features from the 8 calculated features, all 29 points of the waveform, and all 37 total attributes were considered (Appendix E Line # 36-64).

K-means clustering was performed, varying the number of times the algorithm is run with different centroid seeds (10 & 25) and the maximum number of iterations of the algorithm for a single run, from 300 to 700 with a step size of 200. (Appendix E Line # 65-93).

Hyperparameter tuning for t-SNE included varying perplexity, learning rate, and the maximum optimization iterations. The algorithm was run for perplexity values of 25, 50, 100, and 200; learning rate values of 200, 500, and 750; and maximum optimization iterations of 300, 500, and 1000 (Appendix E Line # 94-127).

K-nearest neighbors consisted of varying the number of neighbors from 3, to 5, to 10 (Appendix E Line # 128-154). In gradient tree boosting, 100, 500, and 1000 decision trees were varied, while the learning rate was kept constant at 0.1 (Appendix E Line # 155-181). For the extra trees algorithm, the number of trees was varied from 100, to 500, to 1000 trees (Appendix E Line # 182-208).

Finally, in the logistic regression classification, regularization strength (C) was varied from 1, to 1.5, to 2 (Appendix E Line # 209-235). In all previous algorithms but the random forests, 8 calculated features, the 29 waveform points, and all 37 features were varied as well.

### Multi-channel recording

Multi-channel electrophysiological data was a dataset obtained from Dr. Michael Economo of the Janelia Research Campus (27). The PT_U and PT_L neurons in this dataset were optogenetically tagged to ensure the validity of their classification in a similar manner to the single-channel recording; thus, the specific types of neurons classified were known, again ensuring the mathematical classification’ s validity. Data preprocessing was performed in MatLab R2018 on the spike waveforms, which were a set of 124 to 256 single points that made a waveform when plotted (See Appendix A.2 and A.3 for full code). Each subset of 32 points was a separate waveform from a specific channel; thus, each plotted waveform of 124-256 points was actually 4-8 spikes on neighboring channels at a time point. Thus, prior to feature extraction, the 32-point waveform with maximum amplitude was extracted from the set of 4-8 waveforms, as it provided the most information about the given neuron (Appendix A.2 Line # 106-120; Appendix A.3 Line # 125-139). Feature extraction was done after the largest waveforms were extracted; 11 features were calculated.

#### Feature extraction & pre-processing

Figure 2 shows the two subtypes of PT neurons, PT_L and PT_U; it is seen that the waveforms look extremely similar; thus, feature extraction is needed to find differences between the cell types mathematically.

**Fig. 2.**
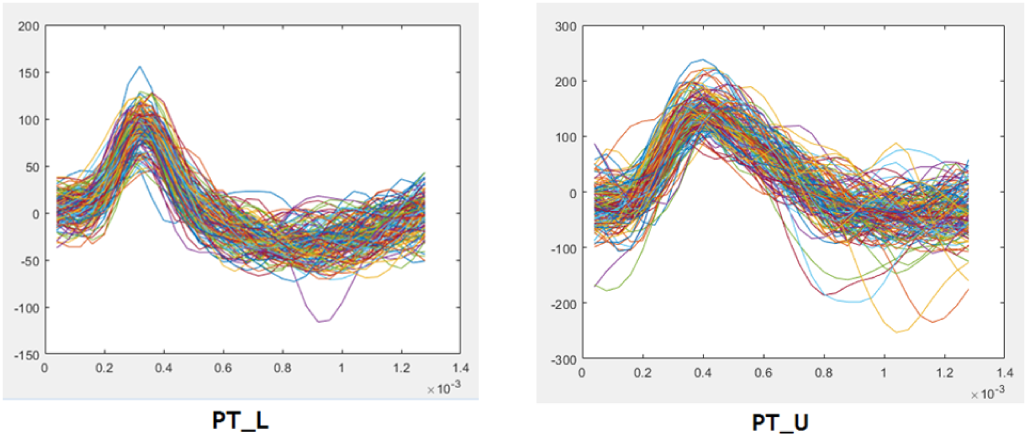
150 waveforms of 29 points superimposed to distinguish the waveform shape between the PT_L & PT_U neuron cell type.

The first eight features were calculated identically to the single-channel dataset, with appropriate adjustments made due to the nature of the provided dataset (Appendix A.2 Line # 122-213; Appendix A.3 Line # 141-232).

The final 3 features were the calculated channel of the neuron, the shank in which it was measured, and the time index at which it was detected (Appendix A.2 Line # 90-96; Appendix A.3 Line # 107-115).

In addition to these 11 calculated features, the entire waveform of 32 units was appended to form the final 32 features. (Appendix A.2 Line # 97-99; Appendix A.3 Line # 116-118).

After this preprocessing, the result was two separate matrices (L & U) with n rows, with n equal to the number of waveforms for a given cell type, and 44 columns, one for each feature (excluding the first, which was the label). These matrices were transferred to a Python IDE (Jupyter Notebook) for further processing and classification via a.csv file intermediary.

#### Training - testing set creation

Further processing was performed in Python 3 to create training and testing arrays in an identical manner to the single-channel recording (See Appendix B for full Python code).

#### MPN & CNN

MPN & CNN were then trained and tested using PyTorch in an identical manner (See Appendix C & D for full Python code of MPN & CNN training and testing).

#### ML algorithms

Seven additional machine learning algorithms were tested identically as well (See Appendix E for full Python code of additional ML algorithms).

## Results

See Appendix F Tables S1 - S3 & Figs. S1 - S18 and Appendix G Tables S4 - S6 & Figs. S19 - S36 for complete data and graphical figures from the MPN, CNN, and other ML algorithms for both single-channel and multi-channel recordings (excluding t-SNE and K-means).

### Single-channel recordings - ANNs

See Figs. S1 - S12 for accuracy and variance plotted as a function of epoch set (1, 2, 3, 5, 10, 25, 50, & 100 epochs). Each figure was initialized to 33% accuracy, to represent a random untrained classifier, since there were three possible classes, along with a maximum variance of 250,000.

In Figures 3 & 4, for the 3 feature subset in both the MPN and CNN, accuracy rose to 70-72% and plateaued after 5 epochs for the MPN and 3 epochs for the CNN. This epochs of convergence is a measure of the speed in which the classifier learned to achieve a consistent accuracy and is indicated by the color of the bars. Variance dropped to nearly 0 as accuracy plateaued for both networks, indicating they were consistently accurate.

**Fig. 3.**
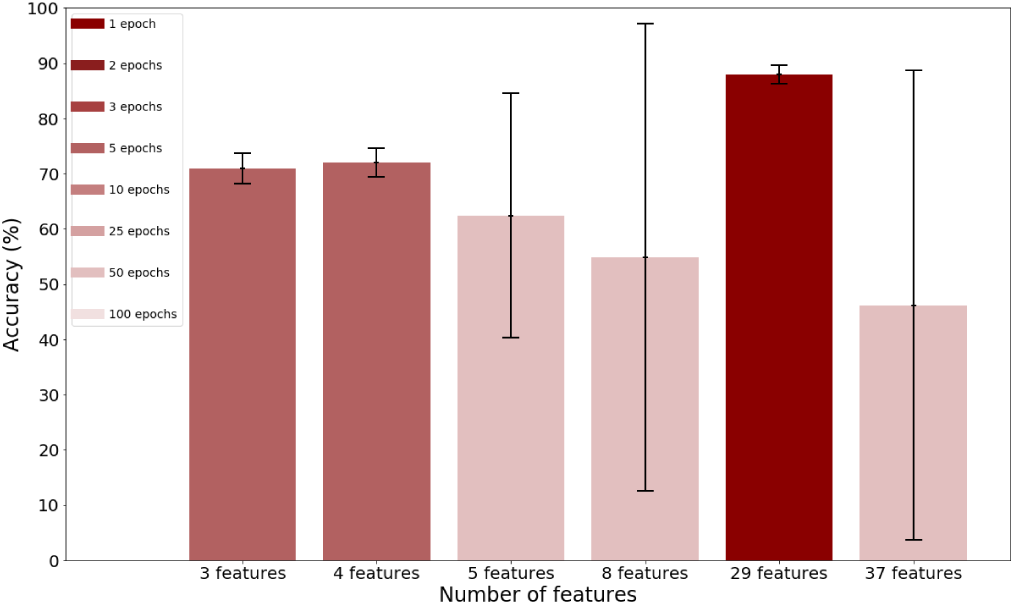
Mean accuracy of MPN as a function of feature subsets, with the black error bars representing variance of the network and the color of the bars represent the epochs of convergence as per the legend

**Fig. 4.**
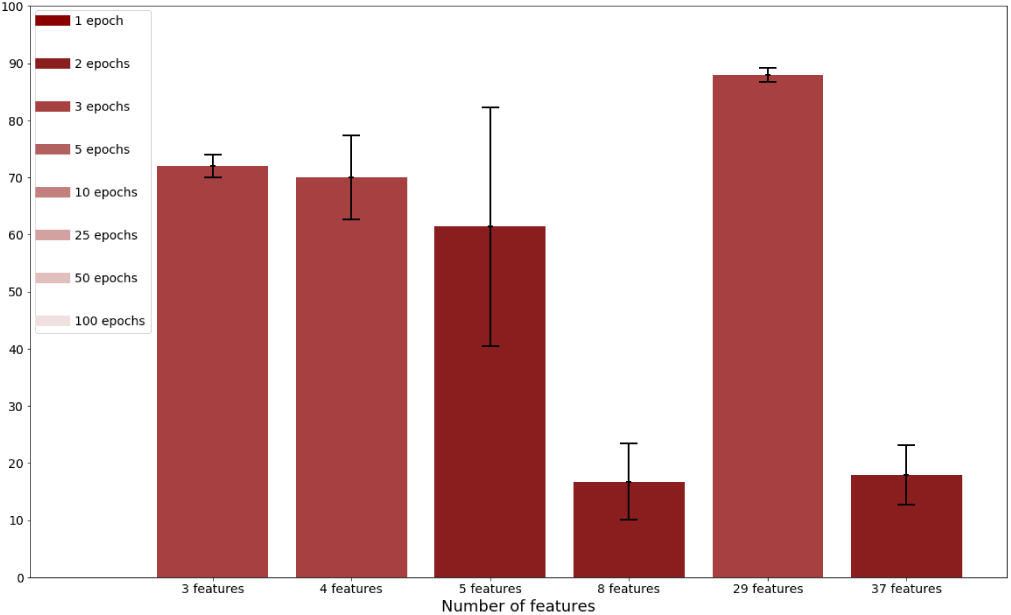
Mean accuracy of the CNN as a function of feature subsets, with the black error bars representing variance of the network and the color of the bars represent the epochs of convergence as per the legend

For the 4 feature subset, accuracy and variance behaved nearly identically, plateauing after the same number of epochs for both the MPN and CNN. For 5 features,accuracy initially rose but then dropped off as epochs reached 100, and variance shot up to 200,000 for both networks. Even when accuracy was at its highest and variance was it its lowest, indicating some reliability at 3 epochs, the accuracy did not exceed 67-73%.

In the 8 feature subset, the networks behaved very differently. Figure 3 shows the MPN, with an accuracy of 55% but extremely high variance indicated by the error bars. Thus, it was unreliable despite decent accuracy. In contrast,Figure 4 shows the CNN with an incredibly low accuracy of 16%, worse than random guessing, even as variance was low; therefore, the CNN was consistently poor. While the networks did behave differently, they provided similar results: the trend towards poorer performance as more calculated features are added suggests the features become increasingly ambiguous between cell types.

Figures 3 & 4 show promising results from both the MPN and CNN with 29 features. With convergence at just 3 epochs, both networks shot up to 88-89% as variance dropped to nearly 0.

The networks behaved similarly for all 37 attributes as they did for the 8 calculated features; for the MPN, accuracy was 46%, while variance remained extremely high. For the CNN, accuracy again dropped below random guessing to 18% with low variance. This poor performance as the eight calculated features were added again indicates the feature calculation was not rigorous, or were potentially not selected ideally. This is especially possible in waveforms with different, unique spike shapes that may have thrown off the feature extraction.

### Single-channel recordings - ML algorithms

See Figs. S13 S18 for detailed graphics of performance as a function of the variable hyperparameters.

#### Random forests

Random forests performed extremely well in classifying the FS, PT, and IT waveforms (see Figure 5). For 3 maximum features, a mean accuracy of 90.5% was seen, which decreased as more calculated features were added; 4 features yielded 89.1%, 5 features yielded 88.1%, and 8 features yielded 87.5%. When all 29 points of the waveform were used by the random forest, a mean accuracy of 98.1% was achieved, which decreased to 95.0% when all 37 attributes were used. Finally, the RF algorithm achieved maximum accuracy at 100 decision trees, with no significant difference between 100, 500, and 1000 trees, allowing for more rapid training.

**Fig. 5.**
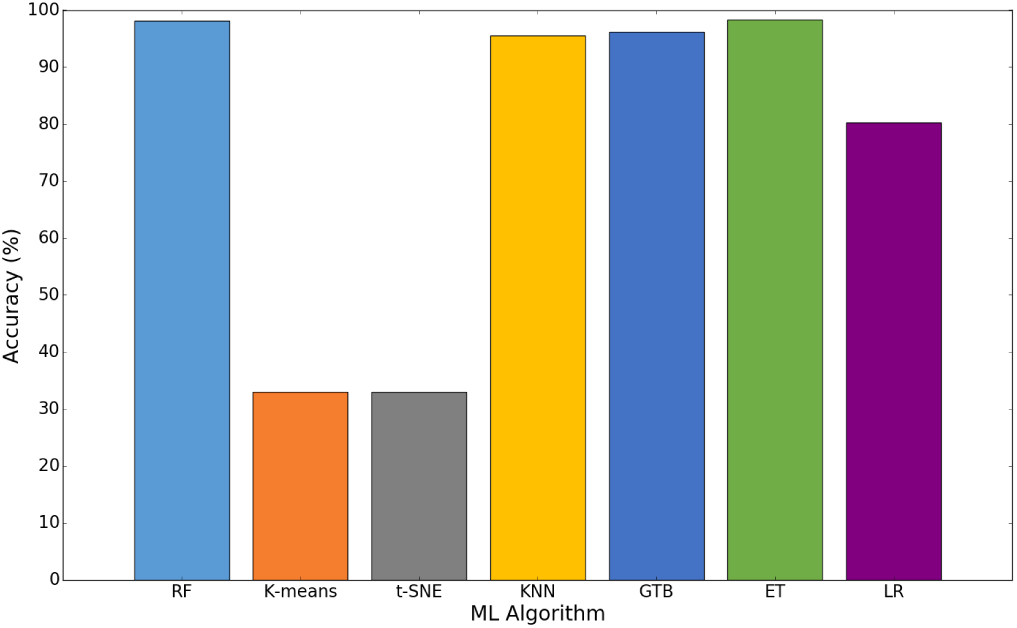
Maximum mean accuracy of various ML algorithms

#### K-means clustering

The k-means clustering was completely inaccurate, yielding accuracies of no greater than random guessing (33%); this occurred for all hyperparameters and feature subsets. It is likely that this poor clustering is because the spikes, though different visually, are difficult to distinguish by the clustering algorithm due to perceived similarities. In addition, there are likely enough anomalies with skewed spike waveforms such that the remaining waveforms cannot be reasonably classified.

#### t-SNE clustering

The t-SNE clustering yielded an image file in which each neuron cell type was assigned a color; green corresponded to FS, blue to PT, and red to IT. Fig. S14 is an example t-SNE clustering output file; all other iterations of t-SNE clustering yielded a nearly identical cluster. The clustering shows that t-SNE is essentially random (33%), likely for a similar reason as K-means clustering.

#### K-nearest neighbors

K-nearest neighbors performed extremely well in classifying the FS, PT, and IT waveforms (see Figure 5). For 8 features, a mean accuracy of 83.8% was seen. When all 29 points of the raw waveform were used by the KNN algorithm, a mean accuracy of 96.2% was achieved, which then decreased back to 83.9% when all 37 attributes were used. The number of neighbors in the KNN algorithm was varied, showing some difference between the 3, 5, and 10 neighbors and indicating that 3 neighbors worked best, since there were 3 classes for the neurons in the ground-truth data.

#### Gradient tree boosting

Gradient tree boosting performed fairly well in classifying the waveforms (see Figure 5). For 8 features, a mean accuracy of 87.8% was seen. When all 29 points of the waveform were used by the GTB algorithm, the mean accuracy decreased to 85.9%, which followed an opposite pattern from the previous algorithms. Interestingly, with all 37 attributes, the mean accuracy shot up to 95.5%, revealing a different set of results than previously seen. The variation in the number of trees in the GTB algorithm showed some difference between the 100, 500, and 1000 trees and indicated that 100 trees worked best.

#### Extra trees

The extra trees classifier performed the best in classifying the waveforms (see Figure 5). For 8 features, a mean accuracy of 92.1% was seen. When all 29 points of the waveform were used by the algorithm, a mean accuracy of 98.3% was achieved, which then decreased slightly to 97.0% when all 37 attributes were used. Although the number of trees in the classifier was varied, there was no significant difference; 100 trees is ideal.

#### Logistic regression

Finally, logistic regression performed the worst of the non-clustering algorithms in classifying the waveforms, exhibiting a maximum accuracy of 83.0% with 29 features. Although the inverse regularization strength in the classifier was varied, there was no significant difference between 1, 1.5, and 2.

Of the ML algorithms tested, RF, KNN, GTB, and ET classification yielded an accuracy of >95% compared to the neural networks, which plateaued at 88-91%. In addition, excluding the GTB algorithm, the 29 points of the raw waveform worked best for all other ML algorithms, while the calculated features threw off the classifiers.

### Multi-channel recordings - ANNs

The classification in the multi-channel recordings was significantly more difficult than in single-channel recordings, due to the similarity between the two waveforms in question (Figure 2). However, effective classification with reasonable accuracy of these neuron cell types is invaluable in large-scale classification, as it would allow for the classification of neuron subtypes within a specific neuron type, to provide a better picture of neural circuitry and more specific relationships between neurons and behavior. See Figs. S19 - S30 for accuracy and variance plotted as a function of epoch set (1, 2, 3, 5, 10, 25, 50, & 100 epochs). Each figure is initialized to 50% accuracy to represent a random untrained classifier, since there were two possible classes, with a maximum variance of 70,000.

Interestingly, seen in Figures 6 and 7, the ANNs did not perform nearly as well for these recordings; this was likely due to the aforementioned large similarity between the two neuron cell types being classified. The accuracy, despite feature selection or epochs for training, hovered around 50%, never exceeding 68%, which was largely an anomaly. The relatively high and inconsistent variance across feature sets and epochs made any accuracy above 50% unreliable, and the networks took 50-100 epochs to converge, indicating it trained slowly, if at all.

**Fig. 6.**
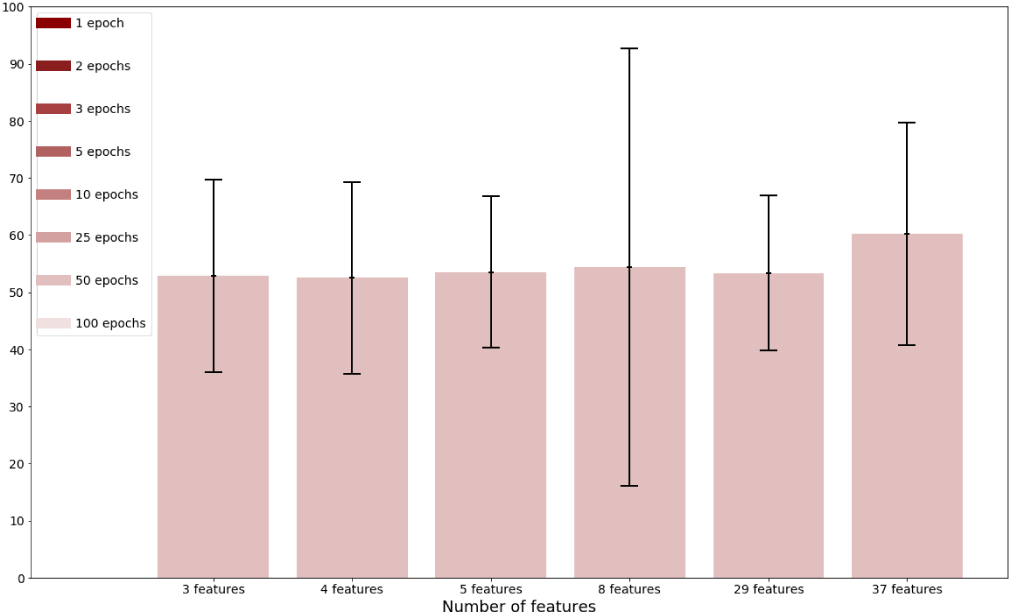
Mean accuracy of MPN as a function of feature subsets, with the black error bars representing variance of the network and the color of the bars represent the epochs of convergence as per the legend

**Fig. 7.**
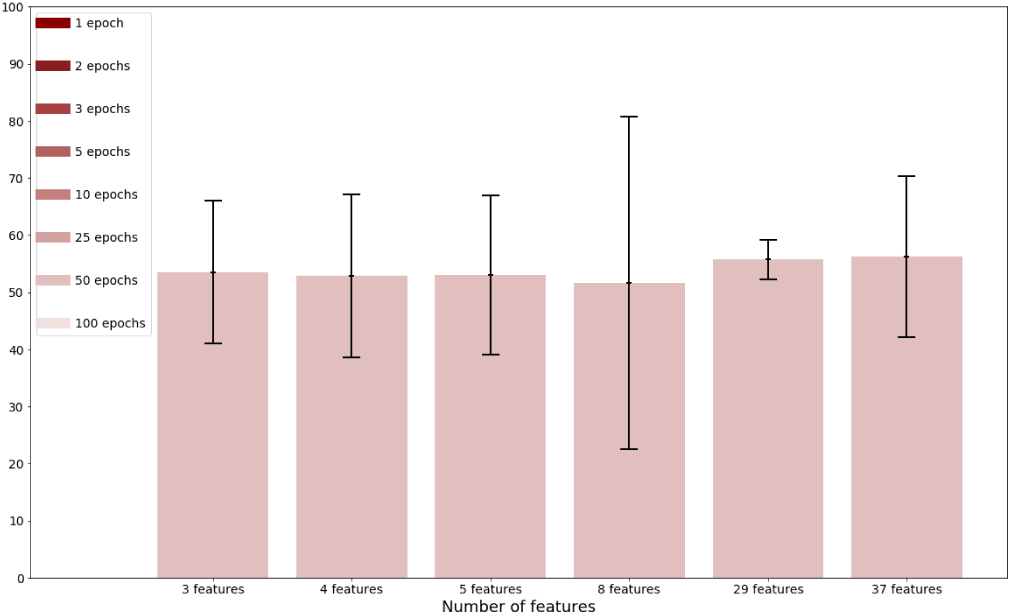
Mean accuracy of CNN as a function of feature subsets, with the black error bars representing variance of the network and the color of the bars represent the epochs of convergence as per the legend

### Multi-channel recordings - ML algorithms

See Figs. S31 - S36 for detailed graphics of performance as a function of the variable hyperparameters.

#### Random forests

Random forests performed fairly well in classifying the PT_L & PT_U waveforms (see Figure 8). For 3 maximum features, a mean accuracy of 75.5% was seen; 4 features yielded 76.1%, 5 features yielded 76.9%, and 8 features yielded 77.2%. When all 32 points of the waveform were used by the random forest, the mean accuracy actually decreased to 73.1%, which then increased to 75.2% with all 43 attributes. Although the number of trees in the random forest algorithm was varied, there was no significant difference between 100, 500, and 1000 trees; again, 100 trees is ideal.

**Fig. 8.**
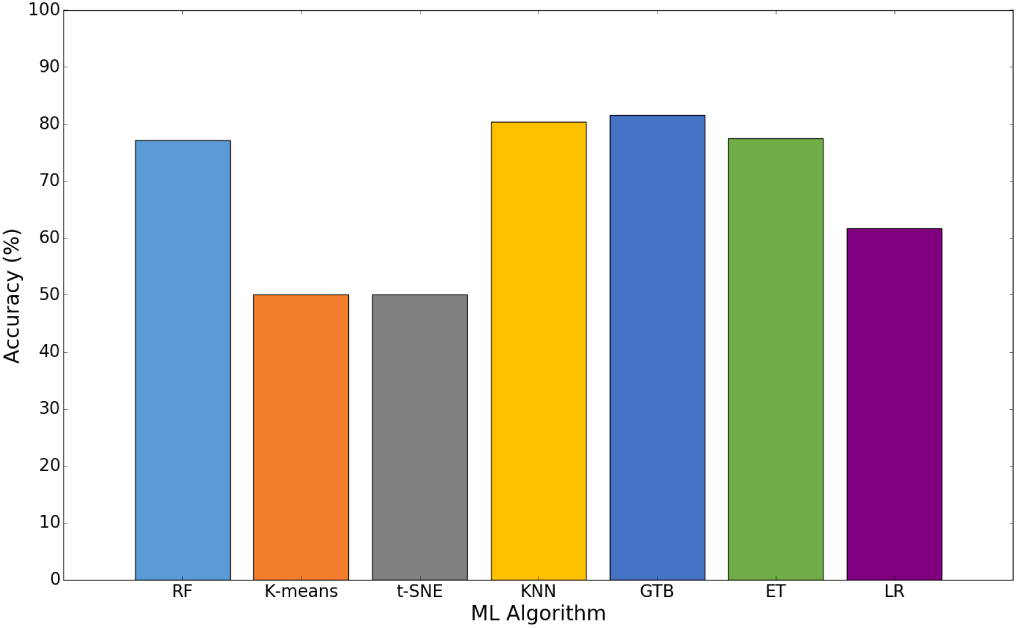
Maximum mean accuracy of various ML algorithms

#### K-means clustering

The k-means clustering was completely inaccurate, yielding accuracies of no greater than random guessing (50%) for all hyperparameters and feature subsets. This is again because the spikes, though different visually, are difficult to distinguish by the clustering algorithm due to some perceived similarities. In addition, there are likely enough anomalies in the dataset with skewed spike waveforms such that the remaining waveforms cannot be reasonably classified.

#### t-SNE clustering

The t-SNE clustering yielded an image file in which each neuron cell type was assigned a color; green corresponded to PT_L, while red corresponded to PT_U. Fig. 32 is an example t-SNE clustering output file; all other iterations of t-SNE clustering with different feature selection and other hyperparameter tuning yielded a nearly identical cluster as shown. The clustering shows that t-SNE is random (33%), and cannot be effectively used to classify these neuron cell types, likely for a similar reason as K-means clustering.

#### K-nearest neighbors

K-nearest neighbors performed the best in classifying the waveforms (see Figure 8). For 8 features, a mean accuracy of 81.6% was seen. When all 32 points of the raw waveform were used by the algorithm, a mean accuracy of 68.5% was achieved, which then increased to 75.4% when all 43 attributes were used. Although the number of neighbors in the KNN algorithm was varied, there was no significant difference between the 3, 5, and 10 neighbors; 3 is ideal since it is closest to the number of classes in the ground-truth data.

#### Gradient tree boosting

Gradient tree boosting performed fairly well in classifying the waveforms. For 8 features, a mean accuracy of 78.4% was seen. When all 32 points of the raw waveform were used by the GTB algorithm, the mean accuracy decreased to 66.7%, and with all 43 attributes, the mean accuracy increased to 73.1%. The number of trees in the GTB algorithm was varied, showing some difference primarily between the 100 trees and 500-1000 trees. In this case, accuracy increased significantly by 2-6% when 500 and 1000 trees; thus, 500 trees are ideal. This increase in optimal decision trees is likely due to the difficulty in classifying the waveforms.

#### Extra trees

The extra trees classifier performed fairly well in classifying the waveforms as well (see Figure 8). For 8 features, a mean accuracy of 77.4% was seen. When all 32 points of the waveform were used by the algorithm, a mean accuracy of 73.5% was achieved, which then increased slightly to 75.4% when all 43 attributes were used. Although the number of trees in the classifier was varied, there was no significant difference between the 100, 500, 1000, & 1500 trees; 100 trees is ideal for maximizing speed.

#### Logistic regression

Finally, logistic regression again performed the worst of the non-clustering algorithms in classifying the waveforms. For 8 features, a mean accuracy of 61.7% was seen. When all 32 raw points of the waveform were used by the algorithm, a mean accuracy of 56.2% was achieved, which then increased slightly to 59.0% when all 43 attributes were used. The inverse regularization strength (IRS) in the classifier was varied, with some difference between the 1, 1.5, and 2; an IRS of 1 performed best.

Of the ML algorithms tested, RF and ET classification yielded an accuracy of >75%, while KNN and GTB performed at >80% accuracy. The ANNs, in comparison, were about 52% accurate on average. In contrast to the single-channel recordings, the 8 calculated features worked best for all ML algorithms, while the raw waveform threw off the classifiers.

## Discussion

In the single-channel recording in which ANNs were used, the initial selection of features (top 3 and 4 features) and all 29 raw waveform points performed most reliably and accurately. This implies that the remaining 4-5 calculated features were not necessarily dependent on cell type, and may change in certain cells. Thus, these features introduced ambiguity to the classifiers, resulting in high variances and mediocre accuracy, or low variances but low accuracy, making the feature subsets unusable for classification. In addition, the highest and most consistent accuracy was seen with the pure waveform alone (88-91% accuracy), indicating that the networks were most adept at selecting and extracting the appropriate features for classification autonomously. This largely eliminates the need for time-consuming data processing and feature extraction.

Training for acceptable accuracy and variance requires no more than 10 epochs for both networks, or 50 epochs for maximum accuracy and minimum variance, both of which can be done within 24-72 hours. After this, running even hundreds of thousands of waveforms through the network, in the case of high density extracellular probes, produces a reliable classification within seconds and can allow for near real-time large-scale classification.

For the single-channel recordings in which the ML algorithms was used, a maximum accuracy of 98% is seen with other ML algorithms. The ideal machine learning algorithm is extra trees due to its consistency and high performance across feature sets; however, random forests, k-nearest neighbors, and gradient tree boosting perform comparably.

For the multi-channel recordings in which the ANNs were used for classification, the high similarity between the neuron subtype waveforms had a large effect on the accuracy. It is seen that ANNs are neither an accurate nor precise method for classifying specific neuron cell subtypes on a large scale, with a maximum accuracy of 68% regardless network architecture. However, when the assorted ML algorithms are applied, a maximum accuracy of 81.6% is seen. The ideal ML algorithm is KNN due to its high performance with the 8 calculated feature set. In addition, gradient tree boosting, random forests, and the extra trees algorithm perform comparably.

The results given above are validated by the use of optogenetic tagging in the datasets that these analyses were performed on, as the true identity of the neurons that were classified was known. Thus, to distinguish between different neuron cell types in the motor cortex, both neural networks and specific machine learning algorithms allow for accurate and consistent classification. In addition, to distinguish between specific neuron cell subtypes in the motor cortex, neural networks are not a viable solution, but specific machine learning algorithms accurately facilitate this large-scale classification.

In comparison to the current broad range of spike sorting methods, many of which are specifically built for high-density electrical probes, ANNs and ML algorithms classify neurons at a much greater rate and with consistently high accuracy. They only require 1 iteration of a bio-imaging technique, while other classification methods require some bio-imaging for every iteration of clustering, to provide ground-truth data for specific neurons in a brain region. Finally, they can accurately classify these neurons without the ambiguity that often results from spike sorting clustering methods. This research reveals a novel, reproducible method for enabling extremely rapid, near realtime large-scale classification at a relatively high accuracy, with widespread applications in all regions of the brain to better understand the connections between neural circuitry and behavior.

## ACKNOWLEDGMENTS

The author received funding from the Janelia Research Campus of the Howard Hughes Medical Institute. The author would like to thank Dr. Louis Scheffer, Dr. Marius Pachitariu, & Dr. Michael Economo for providing the guidance, materials, and expertise needed to conduct this research.

